# Plant Root Expansion Microscopy (ROOT-ExM): A streamlined super resolution method for plants

**DOI:** 10.1101/2024.02.21.581184

**Authors:** Magali S. Grison, Guillaume Maucort, Amandine Dumazel, Dorian Champelovier, Yohann Boutté, Mónica Fernández-Monreal, Emmanuelle M. Bayer

## Abstract

Expansion microscopy (ExM) has revolutionized biological imaging by physically enlarging samples, surpassing the light diffraction limit and enabling nanoscale visualization using standard microscopes. While extensively employed across a wide range of biological samples, its application to plant tissues is sparse. In this work, we present ROOT-ExM, an expansion method suited for stiff and intricate multicellular plant tissues, focusing on the primary root of Arabidopsis thaliana. ROOT-ExM achieves isotropic expansion with a fourfold increase in resolution, enabling super-resolution microscopy comparable to STimulated Emission Depletion (STED) microscopy. Labelling is achieved through immunolocalization, compartment-specific dyes, and native fluorescence preservation, while N-Hydroxysuccinimide (NHS) ester-dye conjugates reveal the ultrastructural context of cells alongside specific labelling. We successfully applied ROOT-ExM to image various cellular structures, including the Golgi apparatus, the endoplasmic reticulum, the cytoskeleton, and wall-embedded structures such as plasmodesmata. When combined with lattice light sheet microscopy (LLSM), ROOT-ExM achieves 3D quantitative analysis of nanoscale cellular process, revealing increased vesicular fusion in close proximity of the cell plate during cell division. Achieving super-resolution fluorescence imaging in plant biology remains a formidable challenge. Our findings underscore that ROOT-ExM provides a remarkable, cost-effective solution to this challenge, paving the way for unprecedented insights into plant cellular subcellular architecture.

**One sentence summary:** ROOT-ExM achieves super-resolution expansion microscopy in plants

## Introduction

Expansion microscopy (ExM) represents a relatively recent addition to the arsenal of super resolution imaging techniques. The primary objective of ExM is to overcome the limitations imposed by the diffraction limit of conventional microscopes by increasing the dimensions of the objects to be observed through isotropic expansion of the biological samples. ExM operates by physically distancing fluorescent probes after fastening them to a gel that can expand. This technique has revolutionized super resolution fluorescence-based microscopy offering a straightforward and cost-effective approach for achieving sub-diffraction resolution limit imaging with standard microscopy setups. Originating from the pioneer work of Edward Boyden in 2015 (Chen et al., 2015), ExM has witnessed extensive advancements in the last decades with the development of protocols designed to accommodate a wide range of biological samples ranging from yeast to zebrafish to neurons or brain tissue (Freifeld et al., 2017; Hinterndorfer et al., 2022; Sarkar et al., 2022) to image proteins, lipids and RNAs (Chen et al., 2015; Chozinski et al., 2016; Ku et al., 2016; Tillberg et al., 2016; Zhao et al., 2017; Gambarotto et al., 2019; Götz et al., 2020; Sun et al., 2021). Remarkably versatile, ExM can be directly applied to pre-labelled samples (immuno- or dye-labelling) or preserve the inherent fluorescence of genetically-tagged proteins. In all case, proteins and antibodies are anchored to the gel by an acrylamide-conjugated molecule such as succinimidyl ester of 6-((acryloyl)amino)hexanoic acid (AcX) or methacrylic acid N-hydroxysuccinimidyl ester (MA-NHS) (Truckenbrodt, 2023). The acrylamide gel containing the water-absorbent sodium acrylate is then digested by Proteinase K, which then allows expansion of the gels with water. The expansion factor is around 4 folds, which theoretically improves the diffraction limited resolution from 200 nm to 50 nm in confocal microscopy. Although expansion microscopy has been swiftly adopted by the scientific community and is conceptually straightforward to implement, it still represents a relatively new technique that is being continuously refined and optimized, when applied to new biological samples with their unique challenges. One domain that has not yet benefited from ExM is, intricate, rigid, multicellular tissues such as those found in plants. In plants, every single cell is encased by a complex network of cellulose microfibrils and cross-linked glycans embedded within a highly interconnected matrix of pectin polysaccharides, which collectively maintain tissue rigidity and shape cells against high internal turgor pressure. Applying ExM to such tissues with inherent internal rigidity is therefore challenging.

In this work, we introduced ROOT-ExM, a modified ExM method designed for the primary root, a fundamental organ for plant growth, nutrient acquisition, pathogen interactions, and responses to the environment (Petricka et al., 2012; Lareen et al., 2016; Ryan et al., 2016; Shekhar et al., 2019; Karlova et al., 2021; Tajima, 2021). The development of ROOT-ExM stemmed from the absence of a robust and comprehensive ExM method suitable for plant tissues. We selected the primary root of *Arabidopsis thaliana* as our focus, given its role as a model system for investigating vital cellular processes, including stem cell niche specification, cell fate determination, cellular polarization, cell division, endocytosis, hormone responses, and numerous other aspects (Jiang et al., 2019; Platre et al., 2019; Retzer et al., 2019; Song et al., 2019; Zhou et al., 2019; Ito et al., 2021; Li et al., 2021; Ruiz-Lopez et al., 2021; Burkart et al., 2022; Gomez et al., 2022; Lebecq et al., 2022).

## Results

### ROOT-ExM physically magnified *Arabidopsis thaliana* primary root by a fourfold factor, with nanoscale isotropy

Adapting ExM to plant organs is primarily challenging due to the presence of cell wall. The wall acts as a physical barrier, hindering antibody penetration (which are relatively large proteins of about 150 kD) and impedes the expansion process because of its rigid polymer network is not affected by the traditional proteinase K digestion step. To implement ExM to *Arabidopsis* root we therefore combined the Protein Retention Expansion Microscopy (ProExM) method (Tillberg et al., 2016) together with a whole-mount immunolocalization protocol commonly employed for *Arabidopsis* root tips (Boutté et al., 2011; Boutté and Grebe, 2014). ProExM involves the direct anchoring of proteins within the target tissue prior to expansion, allowing for the use of common, commercially available fluorescently-labelled antibodies or dyes while also preserving genetically-encoded fluorescent proteins. Whole-mount method achieves immunolocalization and antibody penetration by gently digesting the cell wall with a cocktail of wall-degrading enzymes (Driselase^TM^ consisting of cellulase, endo-1,3-β-glucanase, and xylanase activity), while preserving the cell and root tissue’s integrity. We reasoned that mild-wall digestion will enhance antibody penetration, and simultaneously facilitate tissue expansion.

Combining the two methods we proceeded with the following steps: chemical fixation of the dissected roots, mild wall-digestion, immunolabeling/dye, acrylamide/bisacrylamide incubation, gelation, protein K digestion, and tissue expansion (**Figure 1A)**. We successfully observed antibody penetration and good fluorescent signal before expansion, but we encountered issues with tissue disruption during expansion (ie. non-uniform, anisotropic expansion, internal shearing and tissue rupture). To overcome these technical hurdles, we modified two factors: 1) the cell wall digestion conditions, 2) the gel monomer penetration. After testing various Driselase^TM^ concentrations and incubation times, we determined that a digestion time of 40 minutes at room temperature with a concentration of 2% was optimal for achieving expansion. We also extended the incubation time with the gel monomer solution from 25 minutes to overnight at 4°C, to improve monomer penetration across the tissue. Under these conditions, we successfully achieved expansion of the root tip (from the stem cell niche to the elongation zone of the primary root) (**Figure 1B**). The outer columella cells however resisted expansion and often detached during the process (data not shown).

**Figure 1:**
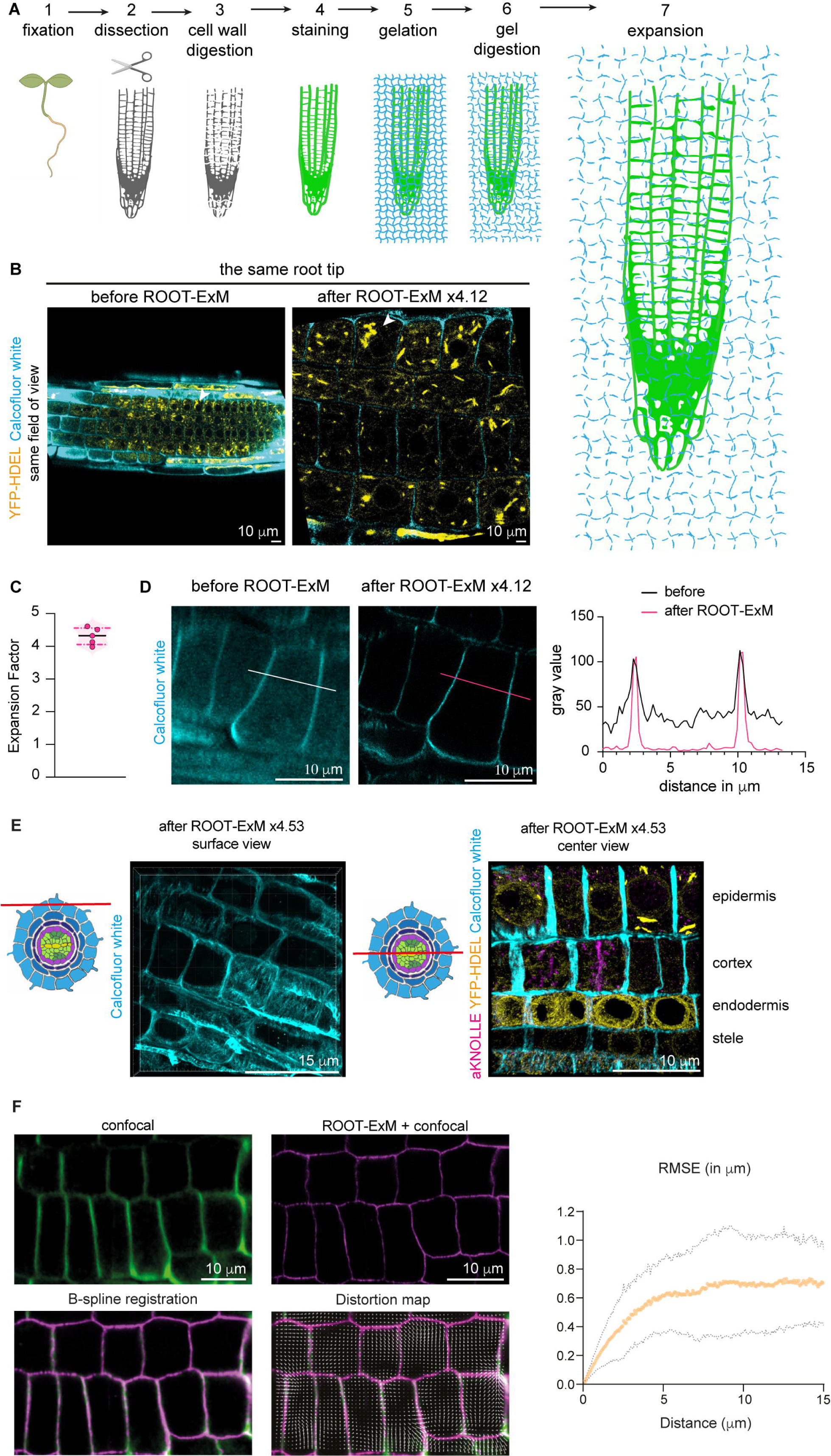
ROOT-ExM achieves fourfold physical and isotropic magnification of *Arabidopsis* root tip. **(A)** Description of the main steps of ROOT-ExM method. **(B)** Confocal images of the same root and field of view before and after ROOT-ExM. Native fluorescence of YFP-HDEL (ER, marker, yellow) expressing *Arabidopsis* root combined with calcofluor white staining (wall marker, cyan). White arrowhead indicates the same cell before and after ROOT-ExM. **(C)** Expansion factor quantification. The diameter of nuclei (stained with DAPI or YFP-HDEL) was measured across five independent experiments (10 nuclei per roots). Mean expansion factor: 4.29 ± 0.26 (mean ± SD). **(D)** Confocal images of the identical area before and after applying ROOT-ExM with Calcofluor white staining of the epidermal walls, along with intensity profiles along red and white lines, illustrating the enhancement in signal-to-noise ratio and resolution. **(E)** Tissue clearing after ROOT-ExM favours in depth imaging. Left panel shows 3D reconstruction of surface signal of cellulose staining with calcofluor white, and right panel shows labelling of the different layers inside the root (Yellow: YFP-HDEL native fluorescence, Cyan: calcofluor white, Magenta: anti-KNOLLE). **(F)** Quantification of the distortion. Representative confocal images of the same field of view before and after ROOT-ExM, with calcofluor white staining of the walls. Lower left panel shows B-spline registration of the images and lower right panel represents the vector field of the registration. Graph shows the root mean square error (RMSE) of four regions from two independent experiments (dashed lines are SD). Scale bar sizes in ROOT-ExM images are calculated by applying the corresponding expansion factor.

We next calculated the expansion factor by measuring the nuclei diameter using DAPI or YFP-HDEL signal (native fluorescence) staining across five independent experiments. We measured a consistent average expansion factor of 4.29 ± 0.26 (mean ± SD; N=5; 10 nuclei per root) (**Figure 1C**). In addition to the physical expansion of the sample, ROOT-ExM also leads to tissue clearing, yielding two key benefits: 1) Enhanced signal-to-noise ratio, exemplified by improved Calcofluor labelling of the walls, allowing finer wall delineation and more accurate thickness measurement (660 nm before ROOT-ExM and 200 nm after ROOT-ExM) (**Figure 1D**). 2) Improved imaging of internal cell layers, due to reduced light scattering in the cleared tissue (**Figure 1E**).

We next tested the accuracy of ROOT-ExM (whether it is isotropic or not) by imaging the same root cells before and after expansion. We analysed the distortions induced by the expansion process by digitally transforming the ROOT-ExM images to fit correlated pre-expansion images and calculated a vector field of deformation (**Figure 1F**). For that, we implemented a user-friendly pipeline to analyse distortions by calculating the expansion errors in correlated images. In this workflow, we first applied image treatment with FIJI on selected regions to facilitate similarity registration (rotation, cropping, scaling). We then performed a rigid registration using the SimpleElastix library (see details in methods) (Chozinski et al., 2016) and extracted the scaling parameter as a correction factor. We then performed a b-spline registration to transform the expanded image for a perfect fit with the original image. This transformation generates a vector field of deformation of the image, and calculates errors between pairs of random points represented by root mean square distances (**Figure 1F**). The distortion analysis showed minimal error distances below 600 nm (root mean square) across length scales up to 15 μm, confirming ROOT-ExM accuracy.

### ROOT-ExM reaches STED super resolution microscopy

The ROOT-ExM expansion factor is approximately fourfold, theoretically enhancing the diffraction-limited resolution by the same factor. This should translate into a gain in resolution and morphological details when imaging subcellular structures. To assess the performance of ROOT-ExM, we selected the NAG1 marker, which labels the medial-Golgi cisterna (Grebe et al., 2003) and compared confocal microscopy, lifetime STED super resolution microscopy (Vicidomini et al., 2018), and ROOT-ExM combined with confocal microscopy. For that, we immunolabeled NAG1-eGFP-expressing *Arabidopsis* lines using nanobodies against GFP. When imaged with confocal, NAG1-labelled Golgi cisterna appeared as small blurry rounded compartments of 1 µm diameter size (**Figure 2A**, left panel). On the contrary, both Lifetime-STED microscopy and ROOT-ExM+confocal microscopy revealed a sharply resolved structure, consisting of a ring composed of small subunits with a thickness ranging from 0.2 to 0.3 µm (**Figure 2A-B**). We therefore concluded that ROOT-ExM is as performant as Lifetime-STED super resolution microscopy imaging.

**Figure 2:**
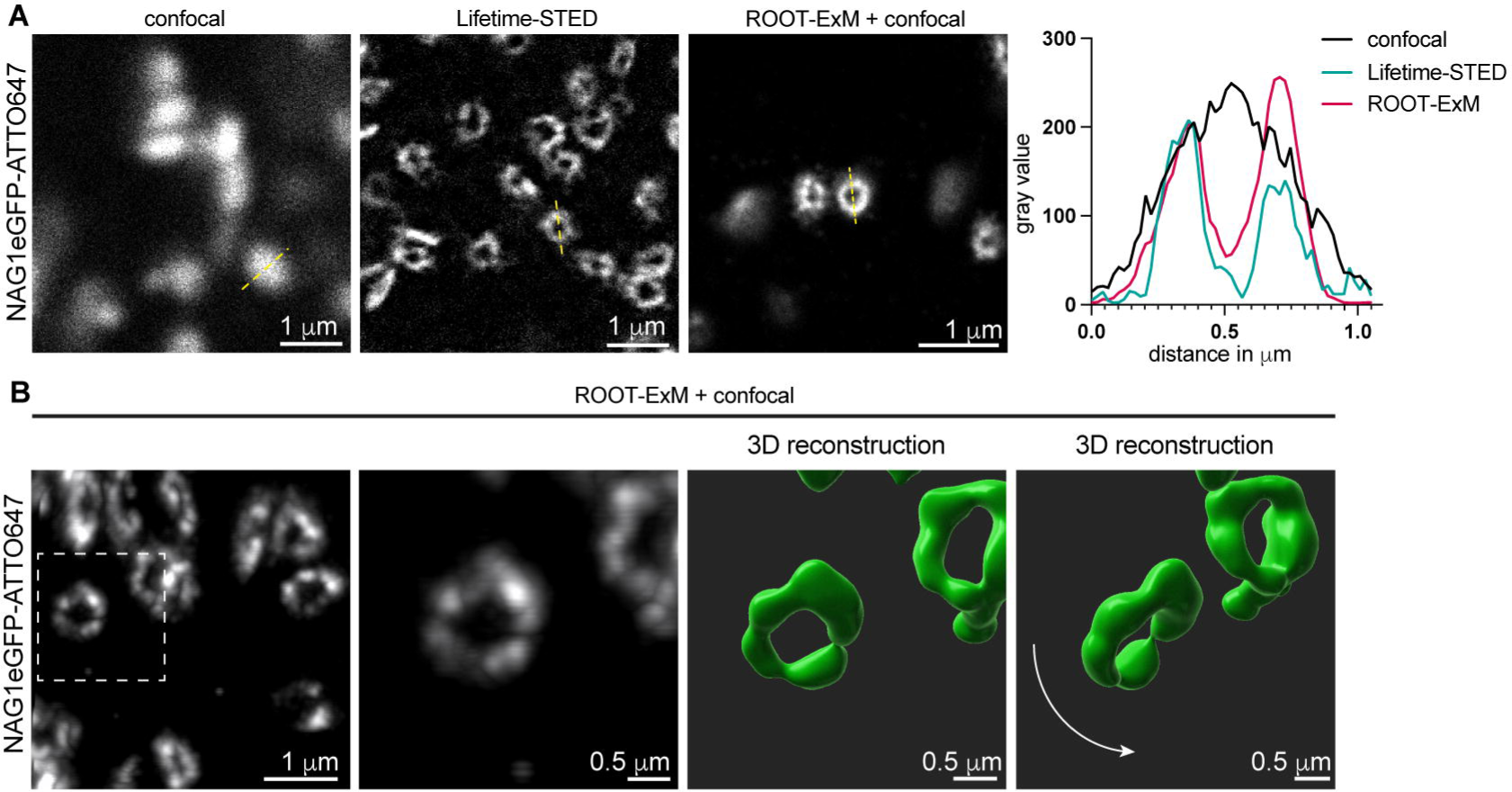
ROOT-ExM allows super resolution imaging using conventional confocal microscope. **(A)** Roots expressing the medial Golgi marker NAG1-eGFP were subjected to immunolabeling with anti-GFP nanobody using either traditional whole-mount immunolocalization (Boutté and Grebe, 2014) or ROOT-ExM. Three representative images of acquisitions in confocal on non-expanded samples (left), super-resolution STED on non-expanded samples (middle) and confocal combined with ROOT-ExM (right). Extreme right panel shows the intensity profile of the yellow dashed lines in the confocal/STED images. **(B)** 3D Imaris reconstruction of medial Golgi cisternae from ROOT-ExM acquisitions revealing the donut shape of the medial cisterna. Scale bar sizes in ROOT-ExM images are calculated by applying the corresponding expansion factor.

### ROOT-ExM is a versatile method that can be applied to cytoskeleton, membrane-, and wall-encased subcellular structures

We then tested the versatility of the ROOT-ExM method by examining its applicability to various sub-cellular compartments. As shown in **Figure 3**, we successfully labelled a variety of cellular structures such as the cell plate (KNOLLE antibody, **Figure 3A-B and K-L**; see also **Figure 5**), the microtubule network (tubulin antibody, **Figure 3C-D**), the cell wall (Calcofluor white dye, **Figure 3 A-D, G-J**), nuclei (DAPI solution, **Figure 3E-F**), plasmodesmata (Callose antibody, **Figure 3G-J**), and the endoplasmic reticulum (ER, YFP-HDEL transgenic lines native fluorescence, **Figure 3K-L**). All tested compartments demonstrated good structural preservation compared to non-expanded samples, and multiple colour labelling was easily achievable (**Figure 1E** and **Figure 3B, D, H, J and L**). ROOT-ExM consistently provided improved resolution for all observed structures compared to non-expanded samples. For example, the labelled microtubules displayed a width of 65 nm ± 1 (mean ± SD) after expansion, contrasting with 334 nm ± 9 (mean ± SD, n=10) before expansion. Altogether, these findings indicate that ROOT-ExM efficiently expands intricate intracellular structures, including the membrane-polysaccharide cell plate compartment during cell division, the ER or the cytoskeleton network, and wall-encased structures like plasmodesmata, to perform super-resolution microscopy with standard fluorophores on diffraction limited microscopes.

**Figure 3:**
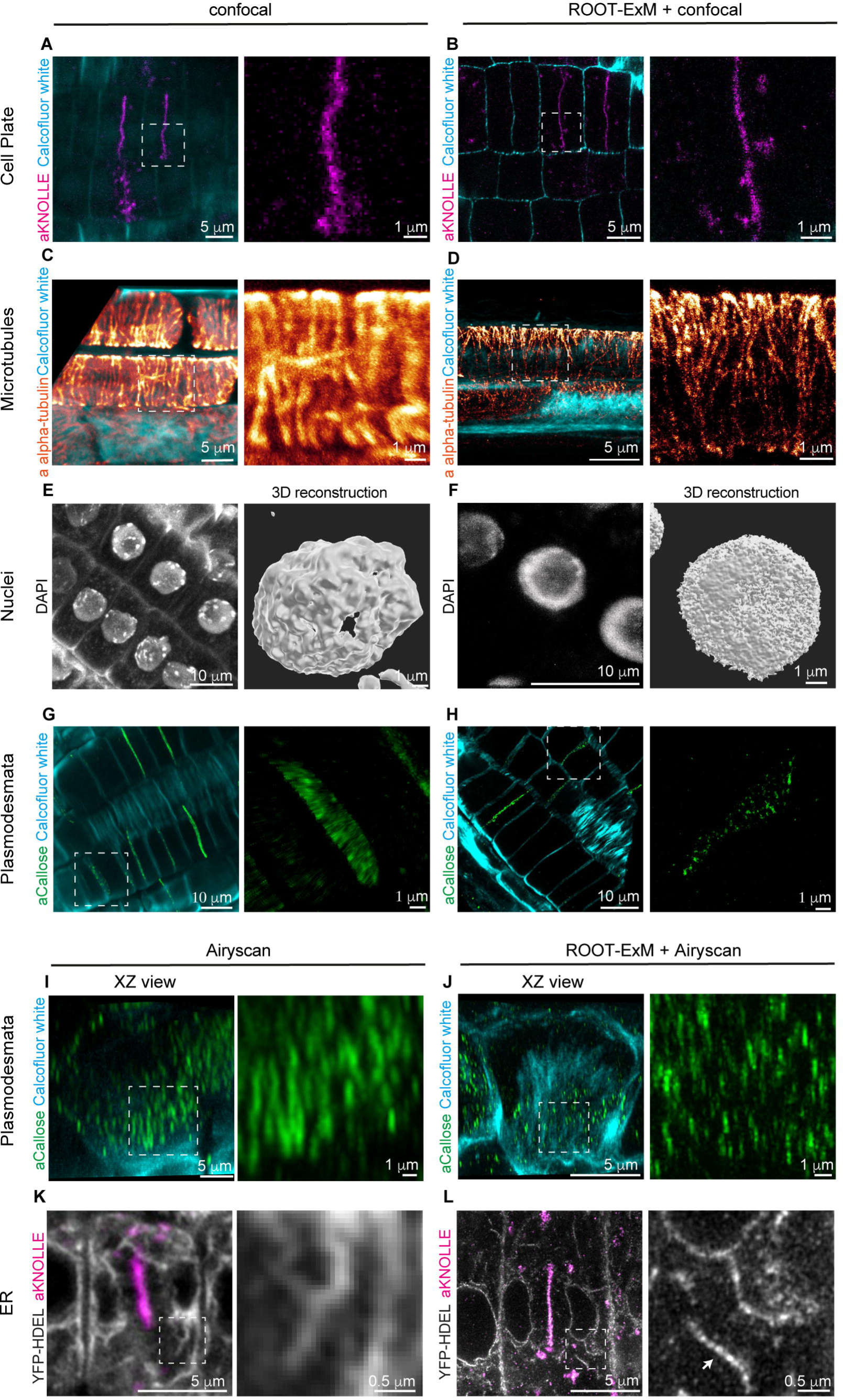
ROOT-ExM enables multicolor imaging of a wide range of subcellular compartments. **(A-L)** Visualization of cellular structures or organelles before (confocal or Airyscan) and after expansion (ROOT-ExM+confocal or ROOT-ExM+Airyscan). (A-B), Cell plate in dividing root epidermal cells visualized using anti-KNOLLE (magenta) immunolabelling. The walls are stained with calcofluor white (cyan). (C-D), Cortical microtubule networks visualized using alpha-tubuline (glow) immunolabelling. The walls are stained with calcofluor white (cyan). (E-F), Nuclei stained with DAPI (grey). (G-J), Plasmodesmata intercellular bridges labelling using an anti-1-3 glucan (callose, green) antibody. The walls are stained with calcofluor white (cyan). (K-L) ER imaging using the native fluorescence of YFP-HDEL (grey) expressing roots. The roots were co-labelled anti-KNOLLE (magenta). Scale bar sizes in ROOT-ExM images are calculated by applying the corresponding expansion factor.

### NHS-labelling combined with ROOT-ExM reveals cellular ultrastructure

We next investigated whether ROOT-ExM can be combined with global protein fluorescent labelling (pan-labelling) to reveal the cellular context of the cells. For this we used N-Hydroxysuccinimide (NHS) ester-dye conjugates, which selectively react with primary amines in proteins and was previously used on animal expanded biological samples (M’Saad and Bewersdorf, 2020). We first tested NHS ester-ATTO488 and NHS ester-ATTO647N. Single cell showed the outline of the nucleus but with little subcellular ultrastructural details (**Figure 4A**). These probes allow non-specific segmentation of nuclei and intracellular organelles by Imaris analysis (**Figure 4A middle panel**). Next, we tried NHS ester-ATTO647 which is compatible for STED microscopy. In our hands, NHS ester-ATTO647 provided stronger and sharper internal signal. Two-dimensional STED revealed cellular ultrastructural details, including hallmark features such as the nuclear envelop, plasma membrane, mitochondria (**Figure 4B**). We were able to spatially resolve details that are normally below diffraction limit, thus invisible to conventional light microscopy such as mitochondria cristae or plasmodesmata intercellular bridges (**Figure 4B-C**). Plasmodesmata intercellular bridges showed an average diameter of 31 nm ± 4 (mean ± SD, n= 6) approaching their actual size as reported in electron microscopy (Nicolas et al. 2017). When using 3D STED microscopy, we could reconstruct the entire volume of these nanoscale intercellular bridges with an isotropic resolution (**Figure 4C**). Finally, NHS-ester labelling is compatible with conventional immuno-fluorescence and chemical staining (**Figure 4D**), therefore allowing for correlative studies that integrate specific and contextual cellular environment.

**Figure 4:**
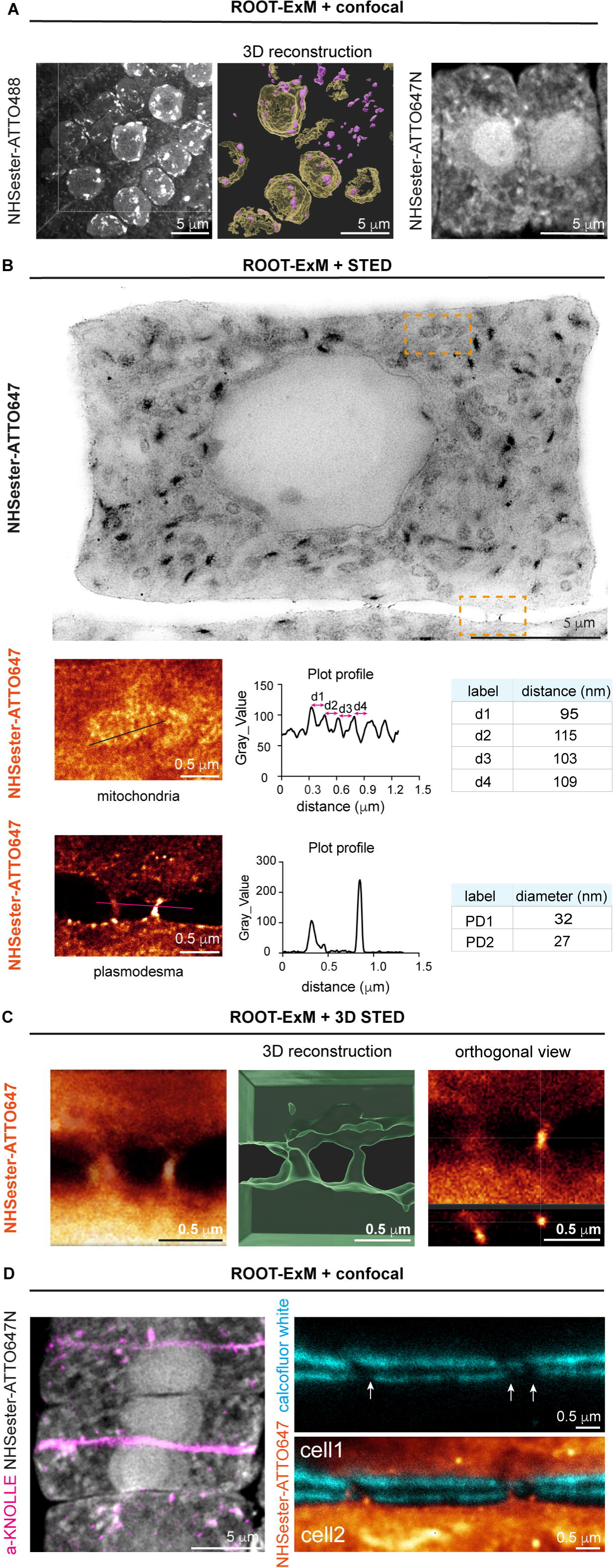
ROOT-ExM combined with NHS-ester pan-labelling allows localisation of proteins in their ultrastructural context. **(A)** NHS-ester ATTO488 and NHS-ester ATTO647N labelling. Middle, Imaris 3D reconstruction and segmentation of NHS-ester ATTO488 labelled nuclei and nucleoles. **(B-C)** STED imaging applied on ROOT-ExM sample labelled with NHS-ester ATTO647. (B) 2D STED. Small inserts show two different compartments identified as mitochondria and plasmodesmata. Intensity profile graphs from the scan lines of the inserts show the resolution of the method. The distance between the mitochondria cristae are approximatively 100 nm while plasmodesmata tube thickness are 27 and 32 nm, very close to their reported diameter in electron microscopy. (C) 3D STED imaging of two plasmodesmata at cell-cell interface from B2. Imaris 3D segmentation of plasmodesmata and orthogonal view showing the isotropic resolution achieved by 3D STED of the plasmodesma channel. (**D**) NHS-ester labelling can be combined with immunolabeling or staining to identify specific structures. (left), NHS-ester ATTO647N labelling combined with KNOLLE (magenta) immunolabeling. (right), NHS-ester ATTO647 labelling combined with calcofluor white staining. Please note that plasmodesmata labelled with NHS-ester ATTO647 are situated in regions characterized by low cellulose content (indicated by arrows and a gap in calcofluor white staining). Scale bar sizes in ROOT-ExM images are calculated by applying the corresponding expansion factor.

### Combining lattice light sheet microscopy and ROOT-ExM for 3D nanoscale quantitative analysis of subcellular processes

To demonstrate the capacity of ROOT-ExM to map 3D subcellular processes that were previously challenging to explore by conventional light-microscopy, we combined lattice light sheet microscopy (LLSM) and ExM. ExM enables super resolution microscopy with a resolution which is similarly improved in the 3 dimensions. In confocal microscopy, the axial (x,y) resolution is typically three times greater than the lateral resolution (z) (see **Figure3 I-J** for instance). This is not the case in light-sheet microscopy where the point spread function (PSF) is less anisotropic (Fernández-Monreal and Ducros, 2021). LLSM also improves the z-resolution by intrinsically sectioning the sample with a 0.5 μm-thin light sheet, allowing rapid imaging at high 3D spatial resolution of large samples with minimal photobleaching (Chen et al., 2014). As a study case, we image the forming cell plate, using KNOLLE as a membrane marker, during cytokinesis (**Figure 5**). The cell plate consists of a membrane-polysaccharide compartment that expands radially to eventually connect to the parental side walls and complete cell division. It is formed by vesicle cargoes that fuse to the equatorial region to form a planar fenestrated sheet that will eventually mature into a cross-wall. When observed through conventional immunolocalization and confocal microscopy on non-expanded samples, the cell plate is seen as a flat, solid structure without internal details (**Figure 5A**). However, combining ROOT-ExM with LLSM reveals the intricate tubulo-vesicular fenestrae membrane structure of the cell plate, allowing us to visualize fenestrae within the otherwise planar cell plate (**Figure 5B, arrowheads**). We obtained 3D images of the cell plate, including cargo vesicles (**Figure 5C**) whose size is below the diffraction limit of optical microscope (Seguí-Simarro et al., 2004). By measuring the size of the vesicles relative to their distance from the cell plate, we noted a significant increase in size as the vesicles approached the middle plane, suggesting localised fusion events (**Figure 5D**). Thus, ROOT-ExM enables quantitative 3D analysis of cellular processes at a nanoscale resolution.

**Figure 5:**
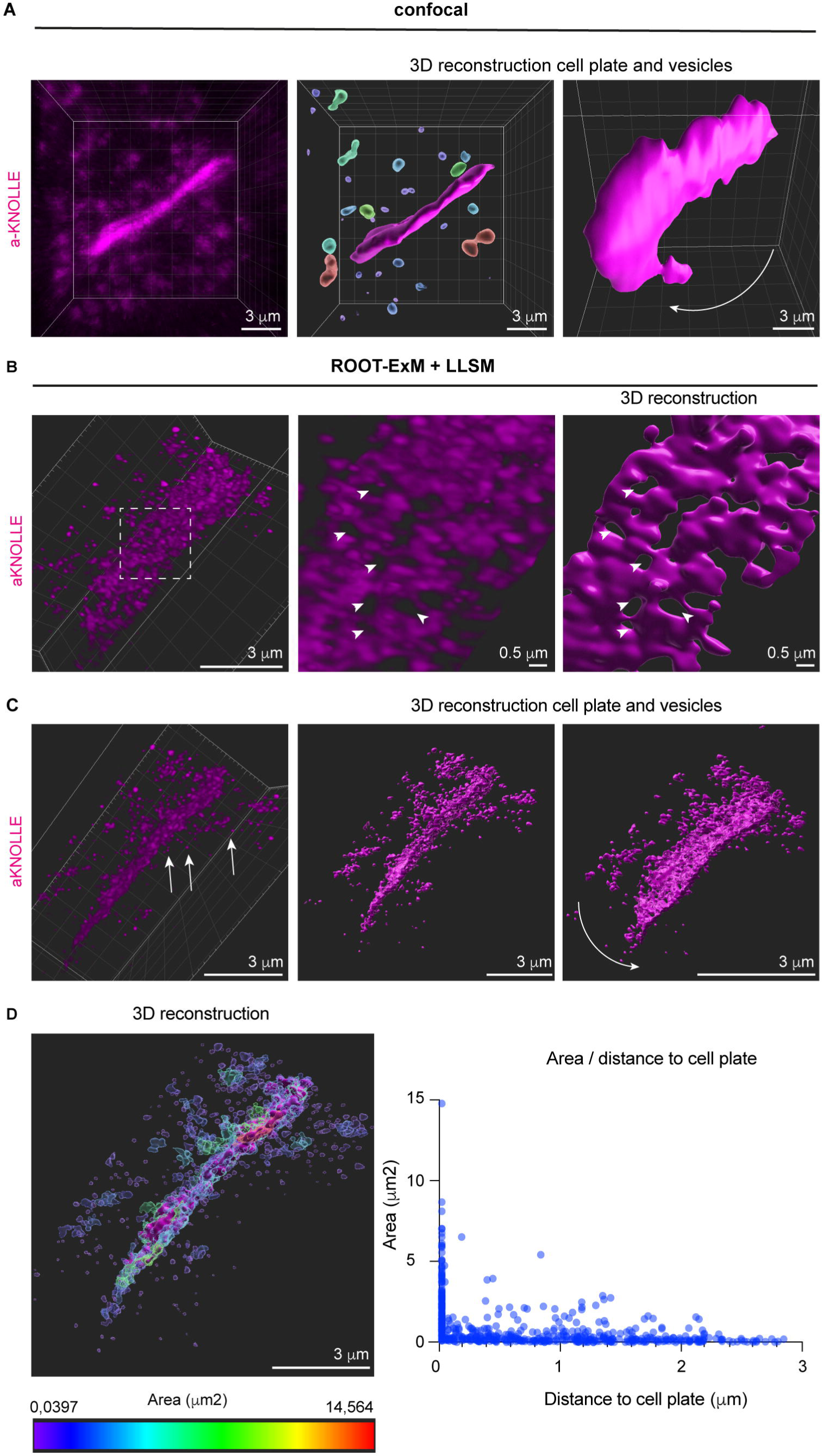
ROOT-ExM combined with lattice light sheet microscopy allow quantitative analysis and 3D isotropic super-resolution. **(A)** *Arabidopsis* roots labelled with KNOLLE antibodies and observed by confocal microscopy prior to ROOT-ExM (left). Imaris 3D reconstruction shows a compact disk-shape cell plate in both lateral (middle) and frontal (right) views. **(B-C)** Volume acquisitions *of Arabidopsis* roots labelled with anti-KNOLLE using ROOT-ExM and imaged by LLSM. (B), Front view of the cell plate showing the intricate tubulo-vesicular structure of the cell plate, including fenestrae (arrowheads). (C) Lateral view of the same cell plate as in (B). Arrows show KNOLLE-labelled vesicles approaching the middle plane. (**D**) Quantification of vesicles size (area) in function of their distance to the cell plate, using Imaris. Scale bar sizes in ROOT-ExM images are calculated by applying the corresponding expansion factor.

## Discussion

In this work we introduce, ROOT-ExM, a robust and straightforward approach for super resolution imaging of plant cells, coupled with a reliable pipeline for image analysis, including expansion factor and distortion assessment.

Development of ExM for plant samples is at its very early stage. One study has reported the use of ExM on isolated chloroplast (Bos et al., 2023) but does not delve into the challenges of expanding intact tissue. Hawkins et al. (Hawkins et al., 2023) developed a protocol tailored for imaging microtubules in liquid-cultured BY2 cells. While their protocol is primarily designed for liquid cultured cells, the authors did attempt root tip expansion using a two-step wall digestion process. However, neither the expansion factor nor tissue distortion were evaluated, nor was the robustness of the protocol assessed. Last, Kao and Nodine applied ExM for imaging complex embryo tissues but with a modest expansion ratio varying from 1.4 to 1.8 (Kao and Nodine, 2021).

Here we offer a reliable method for expanding roots to a factor of 4-folds for super resolution imaging of a wide range of cellular structures. ROOT-ExM boasts another advantage over previous methods as it introduces an analysis workflow that precisely calculates the expansion factor and addresses tissue distortion.

Compared to other super-resolution microscopy techniques ROOT-ExM offers remarkable advantages. Similar to STED microscopy or structured illumination microscopy, ROOT-ExM achieves a fourfold improvement in resolution when compared to confocal microscopy, simply by increasing the size of the biological sample. This enhanced resolution is attained without the need for an expensive and complex specialized super-resolution microscope setup, which often requires a higher level of expertise to operate effectively. For example, ROOT-ExM revealed the distinctive citernae shape of medial Golgi cisternae, achieving a resolution comparable to that of lifetime-STED microscopy. In contrast to many super-resolution techniques, ROOT-ExM provides straightforward access to and enhancement of resolution in three dimensions.

ROOT-ExM is simple to implement. It consists in a sequence of steps involving the incubation of samples with various solutions. The whole root can be processed within four days. ROOT-ExM is not a high throughput method but it is robust and can be done in batch. In our hands we could run of up to 8-10 roots at the same time which is sufficient for conducting an experiment with two different genotypes (gain-of-function, loss-of-function, wild type roots). Imaging can be achieved by epifluorescence, confocal, Airyscan, lattice light sheet microscope within a few hours while data analysis such as the expansion factor or the field of distortion takes from few minutes to few hours depending of the size of the data to analyse. This versatility is valuable because each imaging system possesses distinct advantages, whether it’s related to speed or resolution.

We used the ROOT-ExM to image nanoscale structures within both intracellular compartments and at cell-cell interfaces. These include visualizing the endoplasmic reticulum (by preserving the native YFP-HDEL fluorescence), the Golgi apparatus (using a GFP nanobody against NAG1-eGFP), plasmodesmata intercellular bridges (with the use of primary antibodies against callose and labelling with NHS ester-ATTO647), and the phragmoplast during cell division using a cytokinesis-specific syntaxin KNOLLE (with the use of primary antibodies against KNOLLE). Our findings highlight the versatility of our approach enabling the examination of proteins located within membranes (KNOLLE/NAG1), microtubules as well as structures anchored in the cell wall (callose-labelled plasmodesmata). The fluorescent signal obtained from the secondary synthetic fluorophores is very stable (Alexa Fluor 594, Alexa Fluor 647 and ATTO647N) and although native fluorescence can also be preserved (such as YFP-HDEL), it is also possible to amplify the signal by using anti GFP antibodies (or any other relevant primary antibodies). Please note we were not successful in maintaining tdTomato and RFP native fluorescence.

The ROOT-ExM is also compatible with nuclear DAPI or wall Calcofluor white staining, making the cellular organization within the tissue easily discernible. Additionally, the ROOT-ExM process renders the root completely transparent, similar to clearing techniques. This transparency allows for deep imaging of larger tissue volumes using light microscopy methods, which are often limited by light scattering within the tissue.

Another notable advantage of ROOT-ExM is its ability to provide subcellular context and ultrastructural details, in addition to specific protein localization. This is achieved by employing the cost-effective NHS ester dye-conjugates for staining total protein content. Similar to electron microscopy, which utilizes non-specific stains for lipids and proteins, NHS ester staining in ROOT-ExM allows for the labelling of multiple structures within a cell and yields valuable insights into subcellular organisation at nanoscale resolution. Notably, using NHS ester-ATTO647, we successfully visualized plasmodesmata membrane bridges, which typically have dimensions, no larger than 30 nm in diameter and 100-300 nm in length at cell-to-cell interfaces. By combining pan-labelling with immunodetection, one can confirm the identification of intracellular structures using a specific NHS-ester conjugate. This allows for its direct use as a specific marker without requiring additional permeabilization or incubations. For example, NHS-ester ATTO647 effectively labels plasmodesmata, which are identified by gaps in calcofluor white staining.

We further show that combining ROOT-ExM with LLSM provides a quantitative multicolour 3D super-resolution method. Light sheet fluorescence microscopy is a good alternative to laser scanning approaches in cleared samples since it increases acquisition speed and reduces photobleaching. Among all light sheet fluorescence microscopy techniques, LLSM presents the unique advantage of combining submicrometric spatial resolution over a large field of view, thanks to specific lattice beam shaping (Chen et al., 2014; Fernández-Monreal and Ducros, 2021). As an example of its performance, LLSM reduces the acquisition time of a 20 µm^3^ volume by around 1000-fold compared to confocal laser scanning, with lateral and axial resolutions of 0.3 µm and 0.5 µm, respectively, along with an increased signal-to-noise ratio. The near-isotropic resolution of LLSM allows accurate 3D analysis, and its combination with ExM has already been successfully utilized in thick samples such as mouse cortex and Drosophila brain with excellent outcomes (Gao et al., 2019). Thus, combining ROOT-ExM with LLSM provides a fast, efficient, quantitative, multicolor 3D super-resolution method. In conclusion, we believe that ROOT-ExM will become a widely used super resolution method for nanoscale imaging of intricate intracellular structures with standard fluorophores on diffraction limited microscopes. It will offer unique opportunities for scientists irrespective of their equipment resources to interrogate complex subcellular processes during biotic/abiotic stresses and regulation of root growth and development in wild-type and various mutant backgrounds.

## Methods

### Plant material

Plants were grown 5 days vertically in long day conditions (150 mE/m2/s -1, 16h light/ 8h dark) at 22°C on ½ Murashige and Skoog (Duchefa) media with 1% (w/v) sucrose, 2.5mM morpholinoethanesulfonic acid (Sigma), 8% plant agar, pH5.8 with KOH. Arabidopsis wild type and transgenic lines expressing 35S::YFP-HDEL(Pain et al., 2019) and 35S::NAG1-eGFP were used (Grebe et al., 2003).

### Immunolabeling

The immunolabeling procedure was adapted from Boutté and Grebe, 2014. After seedling fixation (4% paraformaldehyde dissolved in MTSB: 50mM PIPES, 5mM EGTA, 5mM MgSO4 pH 7 with KOH) and washing, the root tips were dissected in excess of water on a glass slide using a razorblade and positioned in the centre of a Corning® BioCoat™ Poly-D-Lysine Glass Coverslip 12 mm (VWR) in 5 µL of water. After the complete dry of the root tip (approximatively 1h) the coverslips were transferred in a 12 wells plate. The immunolabelling procedure is the same as describe in Boutté and Grebe (Boutté and Grebe, 2014), except for the cell wall digestion incubation time that was set up to 40 minutes to optimize expansion. The following antibodies and nanobodies were used: anti-KNOLLE 1/8000 (Boutté et al., 2010), anti-Beta1_3D-Glucan 1/500 (BioSupplies Australia), donkey anti-rabbit-AlexaFluor AF594 1/500 (Abcam), donkey anti-rabbit-ATTO647N 1/500 (Abcam), donkey anti-mouse-AlexaFluor AF594 1/500, donkey anti-mouse-ATTO647N 1/500, GFPbooster-ATTO647N 1/300 (Proteintech).

### Expansion

The proExM method was adapted to plant tissue from Tillberg et al., 2016. After immunolabeling the coverslips were incubated in Acryloyl-X-SE 0.1 mg/mL in PBS (ThermoFischer) overnight at RT and then in the monomer solution (11.7% NaCl, 3 % acrylamide (Sigma), 0.15% N,N′-methylenebisacrylamide (Sigma), 8.625% sodium acrylate (Interchim) and 0.5% 4-Hydroxy-Tempo (Sigma)) for 6 hours at RT and then overnight at 4°C. The coverslips were pre-incubated in the gelling solution (0.2% ammonium persulfate and, 0.2% Temed in 1 mL of monomer solution) for 30 minutes on ice under gentle agitation. Coverslips are then transferred upside down onto a drop of 35 mL of freshly prepared gelling solution in a humid chamber with parafilm, onto a drop of 35 µL of freshly prepared gelling solution, and incubated 2h at 37°C. After gelation, the embedded samples were incubated in digesting solution (50 mM Tris (pH=8.0), 2.5 mM EDTA, 0.5% Triton X-100, 0.8 M Guanidine-HCl and 8 units/mL Proteinase K) at RT overnight. Gels containing the root tip were carefully detached from the coverslip, dissected to eliminate excess of empty gel, transferred into a 6 wells plate and expanded in excess of ddH20. The ddH2O was renewed every 45 minutes until plateau is reached.

### N-Hydroxysuccinimide-ester (NHS-ester) staining

The gels were incubated 1.5h at RT under gentle shaking with NHS-ester ATTO488 (Sigma, 41698), NHS-ester ATTO647 (Sigma, 07376) or NHS-ester ATTO647N (Sigma, 18373) diluted at 20 μg/mL in 100 mM sodium bicarbonate at pH=8.0, then washed 3 times with PBS and expanded. Note that NHS-ester incubation can be done together with calcofluor white and DAPI.

### Confocal Imaging

Distorsion analysis experiments are performed on the same regions of gels before and after expansion. The labelled roots and the expanded gels were mounted on a Corning® BioCoat™ Poly-D-Lysine Glass Coverslip, covered with water and installed in an open chamber dedicated to observation in an upright microscope. Acquisitions were done in sequential mode using a Zeiss LSM 880 equipped with a 40x NA1.2 water dipping objective. Z-stacks series were acquired with a z-step of 0.3-0.5 um. The laser lines used were: 405 nm laser for calcofluor white, the 514 nm line for YFP, and the 594 nm and 630 nm lines for Alexa Fluor^TM^594 and ATTO647N, respectively. The emission filter applied were: 420-460 nm for calcofluor white, 530-570 nm for YFP, 610-640 nm for Alexa Fluor^TM^594 and 655-690 nm for Atto647N.

In some high-resolution images, we used the Airyscan module of the Zeiss LMS880. The configuration of Airy scan acquisitions is: Main Beam Splitter (MBS) 488/561/633 and emission filters BP495-550 + LP570 to observe YFP fluorescence.

### STimulated Emission Depletion (STED) microscopy

STED acquisitions were performed using an inverted LEICA SP8 (DMI6000 TCS SP8 X) equipped with a STED and a FALCON module, and a 93x NA1.3 Glycerol objective. The excitation laser line was 594 nm for the AF594. The power of the excitation laser was set between 5-15%, the power of the 775 nm depletion laser was set to 40%. Emission signal was collected between 610-640 nm with a frame accumulation of 12 for FLIM measurements. Images were automatically processed by the τ-STED module of Leica.

The acquisitions in Figure 5 were done using an Abberior Facility Line (https://abberior.rocks/superresolution-confocal-systems/facility/) equipped with a 60x NA1.2 water objective, a 775 nm pulsed depletion laser, a 2D and 3D STED module, and adaptive optics for image restoration (. We used the 405 and 640 nm laser lines to image calcofluor white and NHS-ester ATTO647, respectively.

### Lattice Light Sheet Microscopy (LLSM)

The LLSM was built according to the technical information provided by the group of E. Betzig at Janelia Research Campus, Howard Hughes Medical Institute (HHMI), USA (Chen et al., 2014) upon signature of a research license agreement between Janelia Research Campus and the National Center of Scientific Research (CNRS, France). The excitation objective was 28.6x NA0.66 (EO; Special Optics). Fluorescence was collected with a 25x CFI Apo LWD NA1.1 detection objective (DO; Nikon) and imaged on a camera sCMOS ORCA-Flash4.0 V2 camera (Hamamatsu). The annular mask minimum and maximum numerical apertures were 0.44 and 0.55, respectively. We characterized the LLSM optical resolution using 170-nm-diameter beads and found values near diffraction of 270 nm laterally and 550 nm axially. Illumination power varied depending on the wavelength, sample brightness, labelling intensity, and depth in slice, which ranged from ∼20 to 200 μW spread over the entire width and thickness of the LLSM excitation plane. Samples were mounted in a 5 mm coverslip and imaged in ultrapure water. Acquired volumes were deskewed and deconvolved using LLSpy (https://github.com/tlambert03/LLSpy.git).

### Imaris analyses

3D reconstructions and analysis were done using the Imaris10 software (Oxford Instruments). Prior to Imaris analysis, a median filter (3x3x1) was applied on the confocal images with FIJI. We then created two separated surfaces for the analysis according to their size: the cell plate and intracellular small compartments. These compartments were sorted according to the distance to the cell plate.

### Distorsion analyses

After defining a common field between two correlated acquisitions, we performed rotations and crops over the expanded image to roughly fit the field depicted in the original sample keeping equal x/y ratio with FIJI. We then rescaled the original image by a factor of 4 (estimated expansion factor) and performed the analysis pipeline within Jupyter notebooks (jupyter.org) (ref to the code available).

The steps of the pipeline are:

1. Rigid registration of the two images using the SimpleElastix library (ref https://simpleelastix.github.io). Blurring of the expanded image allows compensation of the rescale effect to detect common features
2. Rigid transformation – translations, rotations and scaling – for an optimal superposition of the two images. The scaling parameter calculated by the registration algorithm gives us a correction factor which is applied to the estimated expansion factor to obtain the real expansion factor (for the field studied). This step also computes the error between couples of matching points.
3. Spline registration to determine the elastic transformations needed for a complete fit. This is achieved using the B-spline registration functions of the library.
4. Complete registration from the distortion parameters applied on the original non-blurred expanded image.

The spline distortion parameters were then extracted and used to map the deformation vector map over a grid. Note that the vectors representation is arbitrary and that this view only offers relative information. Finally random points are selected over the spine of the images and are used as pairs to calculate the root mean squared error between the original image and the expanded one.

The pipeline has been modified from the original code provided by the Voghan team (Chozinski et al., 2016) and is offered as an open source under request.

## Acknowledgements

We thank Mathieu Ducros for his assistance on the LLSM, Sébastien Marais for the help with IMARIS and Christel Poujol for her help with STED. Microscopy was done at the Bordeaux Imaging Center, member of the national infrastructure France-BioImaging supported by the French National Research Agency (ANR-10-INBS-04). This work was supported by the European Research Council (ERC) under the European union’s Horizon 2020 research and innovation program (project 772103-BRIDGING to E.M.B.), the National Agency for Research (Grant ANR PRPC - ANR-21-CE13-0016-01 DIVCON, E.M.B); the French government in the framework of the IdEX Bordeaux University "Investments for the Future" program / GPR Bordeaux Plant Sciences (E.M.B.). Part of the NHS-ester acquisitions were obtained during the international microscopy school MiFoBio2023. We thank Abberior staff members Gero Schlötel and Frédéric Eghiaian for their assistance in the acquisition of the STED images.

## Author contributions

M.S.G, M. F-M and E.M.B conceived and designed the study and experiments. M.S.G., A.D. and M. F-M performed the experiments and analysed data. G.M. performed the distortion analysis. D.C. helped with the confocal acquisitions in NHS-ester samples during the A020 workshop of MiFoBio2023. E.M.B wrote the paper with the help of M.S.G, M.F-M and Y.B.

## Additional information

**Supplemental table 1.**
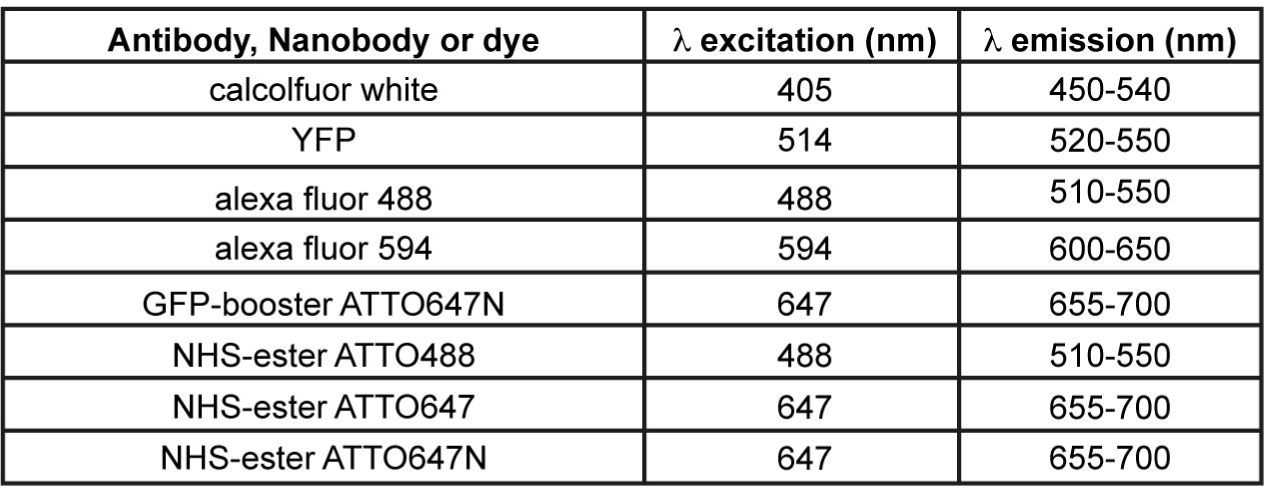

| **Antibody, Nanobody or dye** | **$\lambda$ excitation (nm)** | **$\lambda$ emission (nm)** |
| --- | --- | --- |
| calcofluor white | 405 | 450-540 |
| YFP | 514 | 520-550 |
| alexa fluor 488 | 488 | 510-550 |
| alexa fluor 594 | 594 | 600-650 |
| GFP-booster ATTO647N | 647 | 655-700 |
| NHS-ester ATTO488 | 488 | 510-550 |
| NHS-ester ATTO647 | 647 | 655-700 |
| NHS-ester ATTO647N | 647 | 655-700 |

## BIBLIOGRAPHY

Bos PR, Berentsen J, Wientjes E (2023) Expansion microscopy resolves the 3D thylakoid structure. doi: 10.1101/2023.05.17.541202

Boutté Y, Frescatada-Rosa M, Men S, Chow CM, Ebine K, Gustavsson A, Johansson L, Ueda T, Moore I, Jürgens G, et al (2010) Endocytosis restricts Arabidopsis KNOLLE syntaxin to the cell division plane during late cytokinesis. EMBO J 29: 546–558

Boutté Y, Grebe M (2014) Immunocytochemical fluorescent in situ visualization of proteins in arabidopsis. Methods Mol Biol 1062: 453–472

Boutté Y, Men S, Grebe M (2011) Fluorescent in situ visualization of sterols in Arabidopsis roots. doi: 10.1038/nprot.2011.323

Burkart RC, Strotmann VI, Kirschner GK, Akinci A, Czempik L, Dolata A, Maizel A, Weidtkamp□Peters S, Stahl Y (2022) PLETHORA[WOX5 interaction and subnuclear localization control Arabidopsis root stem cell maintenance. EMBO Rep 23: e54105

Chen BC, Legant WR, Wang K, Shao L, Milkie DE, Davidson MW, Janetopoulos C, Wu XS, Hammer JA, Liu Z, et al (2014) Lattice light-sheet microscopy: Imaging molecules to embryos at high spatiotemporal resolution. Science (80-) 346: 1257998

Chen F, Tillberg PW, Boyden ES (2015) Expansion microscopy. Science (80-) 347: 543–547

Chozinski TJ, Halpern AR, Okawa H, Kim HJ, Tremel GJ, Wong ROL, Vaughan JC (2016) Expansion microscopy with conventional antibodies and fluorescent proteins. Nat Methods 13: 485–488

Fernández-Monreal M, Ducros M (2021) Lattice light sheet microscopy. Imaging Modalities Biol Preclin Res A Compend Vol 1 Part I Ex vivo Biol imaging. doi: 10.1088/978-0-7503-3059-6ch8

Freifeld L, Odstrcil I, Förster D, Ramirez A, Gagnon JA, Randlett O, Costa EK, Asano S, Celiker OT, Gao R, et al (2017) Expansion microscopy of zebrafish for neuroscience and developmental biology studies. Proc Natl Acad Sci U S A 114: E10799–E10808

Gambarotto D, Zwettler FU, Le Guennec M, Schmidt-Cernohorska M, Fortun D, Borgers S, Heine J, Schloetel JG, Reuss M, Unser M, et al (2019) Imaging cellular ultrastructures using expansion microscopy (U-ExM). Nat Methods 16: 71–74

Gao R, Asano SM, Upadhyayula S, Pisarev I, Milkie DE, Liu TL, Singh V, Graves A, Huynh GH, Zhao Y, et al (2019) Cortical column and whole-brain imaging with molecular contrast and nanoscale resolution. Science (80-). doi: 10.1126/science.aau8302

Gomez RE, Chambaud C, Lupette J, Castets J, Pascal S, Brocard L, Noack L, Jaillais Y, Joubès J, Bernard A (2022) Phosphatidylinositol-4-phosphate controls autophagosome formation in Arabidopsis thaliana. Nat Commun. doi: 10.1038/s41467-022-32109-2

Götz R, Kunz TC, Fink J, Solger F, Schlegel J, Seibel J, Kozjak-Pavlovic V, Rudel T, Sauer M (2020) Nanoscale imaging of bacterial infections by sphingolipid expansion microscopy. Nat Commun. doi: 10.1038/s41467-020-19897-1

Grebe M, Xu J, Mö W, Ueda T, Nakano A, Geuze HJ, Rook MB, Scheres B (2003) Arabidopsis sterol endocytosis involves actin-mediated trafficking via ARA6-positive early endosomes. Curr Biol 13: 1378–1387

Hawkins TJ, Robson JL, Cole B, Bush SJ (2023) Expansion Microscopy of Plant Cells (PlantExM). Methods Mol. Biol. Humana Press Inc., pp 127–142

Hinterndorfer K, Laporte MH, Mikus F, Tafur L, Bourgoint C, Prouteau M, Dey G, Loewith R, Guichard P, Hamel V (2022) Ultrastructure expansion microscopy reveals the cellular architecture of budding and fission yeast. J Cell Sci. doi: 10.1242/jcs.260240

Ito Y, Esnay N, Platre MP, Wattelet-Boyer V, Noack LC, Fougère L, Menzel W, Claverol S, Fouillen L, Moreau P, et al (2021) Sphingolipids mediate polar sorting of PIN2 through phosphoinositide consumption at the trans-Golgi network. Nat Commun. doi: 10.1038/s41467-021-24548-0

Jiang Z, Zhou X, Tao M, Yuan F, Liu L, Wu F, Wu X, Xiang Y, Niu Y, Liu F, et al (2019) Plant cell-surface GIPC sphingolipids sense salt to trigger Ca2+ influx. Nature 572: 341–346

Kao P, Nodine MD (2021) Application of expansion microscopy on developing Arabidopsis seeds. Methods Cell Biol. Academic Press Inc., pp 181–195

Karlova R, Boer D, Hayes S, Testerink C (2021) Root plasticity under abiotic stress. Plant Physiol 187: 1057–1070

Ku T, Swaney J, Park JY, Albanese A, Murray E, Hun Cho J, Park YG, Mangena V, Chen J, Chung K (2016) Multiplexed and scalable super-resolution imaging of three-dimensional protein localization in size-adjustable tissues. Nat Biotechnol 34: 973–981

Lareen A, Burton F, Schäfer P (2016) Plant root-microbe communication in shaping root microbiomes. Plant Mol Biol 90: 575–587

Lebecq A, Doumane M, Fangain A, Bayle V, Leong JX, Rozier F, Del Marques-Bueno M, Armengot L, Boisseau R, Simon ML, et al (2022) The Arabidopsis SAC9 enzyme is enriched in a cortical population of early endosomes and restricts PI(4,5)P2 at the plasma membrane. Elife. doi: 10.7554/eLife.73837

Li L, Verstraeten I, Roosjen M, Takahashi K, Rodriguez L, Merrin J, Chen J, Shabala L, Smet W, Ren H, et al (2021) Cell surface and intracellular auxin signalling for H+ fluxes in root growth. Nature 599: 273–277

M’Saad O, Bewersdorf J (2020) Light microscopy of proteins in their ultrastructural context. Nat Commun. doi: 10.1038/s41467-020-17523-8

Pain C, Kriechbaumer V, Kittelmann M, Hawes C, Fricker M (2019) Quantitative analysis of plant ER architecture and dynamics. Nat Commun 10: 1–15

Petricka JJ, Winter CM, Benfey PN (2012) Control of arabidopsis root development. Annu Rev Plant Biol 63: 563–590

Platre MP, Bayle V, Armengot L, Bareille J, del Mar Marquès-Bueno M, Creff A, Maneta-Peyret L, Fiche JB, Nollmann M, Miège C, et al (2019) Developmental control of plant Rho GTPase nano-organization by the lipid phosphatidylserine. Science (80-) 364: 57–62

Retzer K, Akhmanova M, Konstantinova N, Malínská K, Leitner J, Petrášek J, Luschnig C (2019) Brassinosteroid signaling delimits root gravitropism via sorting of the Arabidopsis PIN2 auxin transporter. Nat Commun. doi: 10.1038/s41467-019-13543-1

Ruiz-Lopez N, Pérez-Sancho J, del Valle AE, Haslam RP, Vanneste S, Catalá R, Perea-Resa C, van Damme D, García-Hernández S, Albert A, et al (2021) Synaptotagmins at the endoplasmic reticulum–plasma membrane contact sites maintain diacylglycerol homeostasis during abiotic stress. Plant Cell 33: 2431–2453

Ryan PR, Delhaize E, Watt M, Richardson AE (2016) Plant roots: Understanding structure and function in an ocean of complexity. Ann Bot 118: 555–559

Sarkar D, Kang J, Wassie AT, Schroeder ME, Peng Z, Tarr TB, Tang AH, Niederst ED, Young JZ, Su H, et al (2022) Revealing nanostructures in brain tissue via protein decrowding by iterative expansion microscopy. Nat Biomed Eng 6: 1057–1073

Seguí-Simarro JM, Austin JR, White EA, Staehelin LA (2004) Electron tomographic analysis of somatic cell plate formation in meristematic cells of Arabidopsis preserved by high-pressure freezing. Plant Cell 16: 836–856

Shekhar V, St□ckle D, Thellmann M, Vermeer JEM (2019) The role of plant root systems in evolutionary adaptation. Curr. Top. Dev. Biol. Academic Press Inc., pp 55–80

Song JH, Kwak S, Nam KH, Schiefelbein J, Lee MM (2019) QUIRKY regukates root epidermal cell patterning through stabilizing SCRAMBLED to control CAPRICE movement in Arabidopsis. Nat Commun 10: 1–12

Sun D en, Fan X, Shi Y, Zhang H, Huang Z, Cheng B, Tang Q, Li W, Zhu Y, Bai J, et al (2021) Click-ExM enables expansion microscopy for all biomolecules. Nat Methods 18: 107–113

Tajima R (2021) Importance of individual root traits to understand crop root system in agronomic and environmental contexts. Breed Sci 71: 13–19

Tillberg PW, Chen F, Piatkevich KD, Zhao Y, Yu CC, English BP, Gao L, Martorell A, Suk HJ, Yoshida F, et al (2016) Protein-retention expansion microscopy of cells and tissues labeled using standard fluorescent proteins and antibodies. Nat Biotechnol 34: 987–992

Truckenbrodt S (2023) Expansion Microscopy: Super-Resolution Imaging with Hydrogels. Anal Chem 95: 3–32

Vicidomini G, Bianchini P, Diaspro A (2018) STED super-resolved microscopy. Nat Methods 2018 153 15: 173–182

Zhao Y, Bucur O, Irshad H, Chen F, Weins A, Stancu AL, Oh EY, Distasio M, Torous V, Glass B, et al (2017) Nanoscale imaging of clinical specimens using pathology-optimized expansion microscopy. Nat Biotechnol 35: 757–764

Zhou W, Lozano-Torres JL, Blilou I, Zhang X, Zhai Q, Smant G, Li C, Scheres B (2019) A Jasmonate Signaling Network Activates Root Stem Cells and Promotes Regeneration. Cell 177: 942–956.e14

